# Maternal Under-Nutrition During Pregnancy Alters the Molecular Response to Over-Nutrition in Multiple Organs and Tissues in Juvenile Offspring

**DOI:** 10.1101/2023.07.07.548121

**Authors:** Laura A. Cox, Sobha Puppala, Jeannie Chan, Angelica M. Riojas, Kenneth J. Lange, Shifra Birnbaum, Edward J. Dick, Anthony G. Comuzzie, Mark J. Nijland, Cun Li, Peter W. Nathanielsz, Michael Olivier

## Abstract

Previous studies suggest that mismatch between fetal and postnatal nutrition predisposes individuals to metabolic diseases. We examined whether NHP juvenile offspring of pregnancies with maternal undernutrition (MUN) had altered response to a high-fat, high-carbohydrate diet plus sugar drink challenge (HFCS) compared with controls (CON). Pregnant baboons were fed *ad libitum* (CON) or 30% calorie reduction from 0.16 gestation through lactation, and weaned offspring fed chow diet *ad libitum*. Offspring of MUN were growth restricted at birth. At ∼4.5y offspring received a 7-week HFCS challenge. Baseline and HFCS gene expression in liver, omental fat, and skeletal muscle, liver glycogen content, and fat cell size were quantified. MUN offspring had lower BMI and liver glycogen compared with CON. Pathway analysis showed differences for skeletal muscle and liver, including hepatic splicing and unfolded protein response. MUN offspring consumed more sugar drink than CON. After HFCS, MUN BMIs were similar to CON. Liver showed coordinated response to HFCS in CON but not MUN. Pathway and liver glycogen differences between MUN and CON at baseline indicate *in utero* programming persists in MUN juveniles. MUN catchup growth during HFCS suggests increased risk of obesity, diabetes, and cardiovascular disease. Greater sugar drink consumption in MUN demonstrates altered appetitive drive due to programming. Differences in blood leptin concentrations, omental adipocyte cell size, liver glycogen content, and tissue-specific molecular response to HFCS challenge suggest MUN significantly impacts juvenile offspring ability to manage an energy rich diet.

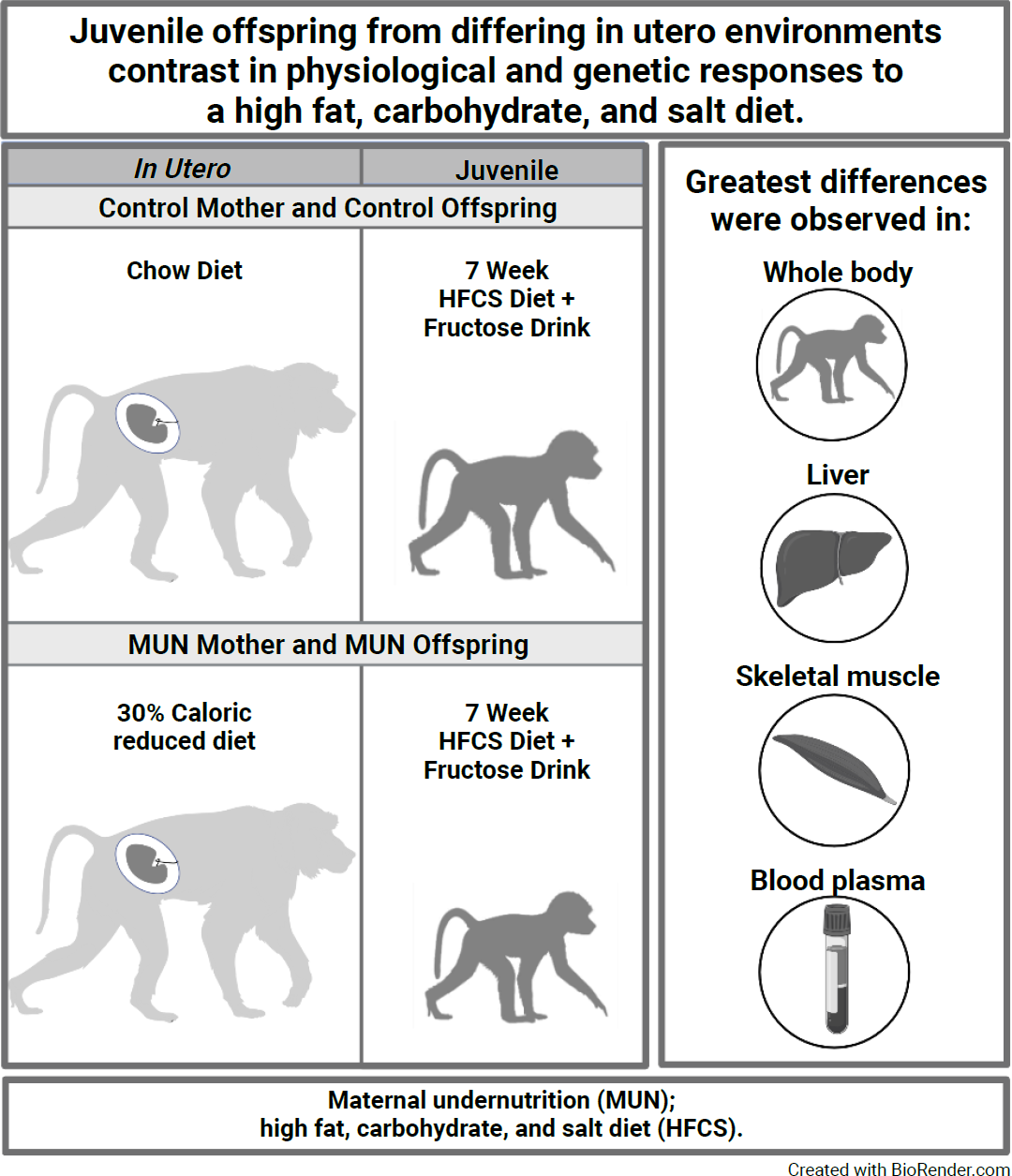

## INTRODUCTION

In 2021, the USDA revealed that 12.5% of US households with women of reproductive years and children experienced food insecurity ^1^. Poor maternal nutrition during pregnancy is linked to intrauterine growth restriction (IUGR) and adverse lifecourse health outcomes in offspring. Human epidemiological studies and controlled nutrient reduction studies in rats and sheep show that a sub-optimal intrauterine environment leading to IUGR alters the trajectory of fetal development with profound effects on life-time offspring health (reviewed in ^2^). IUGR is strongly associated with heart disease, hypertension, obesity, and diabetes in adult offspring of these suboptimal pregnancies. For example, rats fed low-protein diets have high blood pressure from early postnatal life ^3^. In addition, offspring of low-protein fed dams have increased susceptibility to diabetes, insulin resistance, and hypertension when fed a high-fat diet ^4^, similar to what has been observed in human epidemiological studies ^5^.

To date, studies of mechanisms by which maternal under nutrition (MUN) during pregnancy programs offspring health have been conducted primarily in rodents making it difficult to extrapolate to humans due to significant developmental (and nutritional) differences between rodents and nonhuman primates (NHP) ^6^. For example, at birth several rodent organs such as kidney are developmentally similar to mid-gestation stages in primates due to the need for rodent organs to be functional at birth ^7^. This difference alters the trajectory of rodent organ development compared with primates. These differences are apparent for both cellular and molecular properties ^8^. The nonhuman primate (NHP) model of MUN leading to IUGR offers the advantages of defining offspring phenotype and mechanisms using a model that is physiologically and genetically similar to humans and for which the environment can be controlled.

In previous studies, we showed that MUN (70% of total calories fed *ad libitum* control (CON) diet) during pregnancy in NHP results in IUGR in both female and male offspring ^9^. We demonstrated that MUN effects the transcriptome and energy storage in the developing fetal liver via epigenetic changes in PEPCK signaling ^10, 11^. In addition, we found effects on the fetal liver transcriptome through increased RNA splicing and potential metabolic alterations with increased enrichment of genes for fatty acid beta oxidation signaling ^12^. Abundance of orexigenic neuropeptide Y (NPY) was increased, anorexigenic proopiomelanocortin (POMC) was decreased, and leptin signaling downregulated in the hypothalamus of 0.9G IUGR fetuses ^13^, consistent with reduced fetal nutrition programing offspring for increased appetite and later life obesity. These changes are like those shown in rodent models where gene expression for orexigenic peptides increases with corresponding decreases in gene expression for anorexigenic peptides in hypothalamic arcuate (ARH) nuclei of IUGR offspring ^14^. These hypothalamic peptide and leptin feedback signaling changes suggest that MUN resulting in IUGR alters energy sensing pathways and increases appetite. Consistent with these findings, studies in pre-pubertal MUN baboons show altered peripheral insulin sensitivity, potentially predisposing these animals to early onset of type 2 diabetes ^15^.

To explore whether metabolic effects of MUN during fetal development persist in postnatal offspring, we characterized juvenile baboon IUGR offspring aged ∼4.5 years old and age-matched CON that were maintained on a low cholesterol, low fat diet from weaning until study initiation, and examined whole body, cellular, and molecular responses after a 7-week *ad libitum* high-fat, high-carbohydrate, high-salt Western style diet supplemented with a high fructose drink (HFCS). While there are clearly differences between juvenile MUN and CON offspring before challenge, the response to the HFCS challenge emphasizes a dramatic modulation of responses across multiple tissues in MUN offspring compared to CON animals, demonstrating that *in utero* exposure to MUN alters molecular pathways and regulatory mechanisms that likely lead to major changes in appetite, and energy management and storage in IUGR juvenile offspring.

## METHODS

### Ethics Statement

All animal procedures and study protocols were approved by the Institutional Animal Care and Use Committee at Texas Biomedical Research Institute and conducted in Association for Assessment and Accreditation of Laboratory Animal Care approved facilities at the Southwest National Primate Research Center.

### Animal Selection and Management

Baboon housing, feeding, and environmental enrichment were previously published ^16^. All dams spontaneously delivered offspring at full term. Offspring were reared with their mothers in group housing until weaning at ∼9 months, and then maintained on chow diet until diet challenge.

### HFCS Diet Challenge and Sugar Drink Consumption

At ∼4.5 years of age (human equivalent 13 years), 6 CON offspring (3 females, 3 males) and 6 age-matched MUN offspring (3 females, 3 males) were challenged with a 7-week *ad libitum* HFCS diet, high fructose drink with free access to water (Fig. 1A). HFCS diet and high fructose drink details were previously published ^17^.

**Fig. 1:**
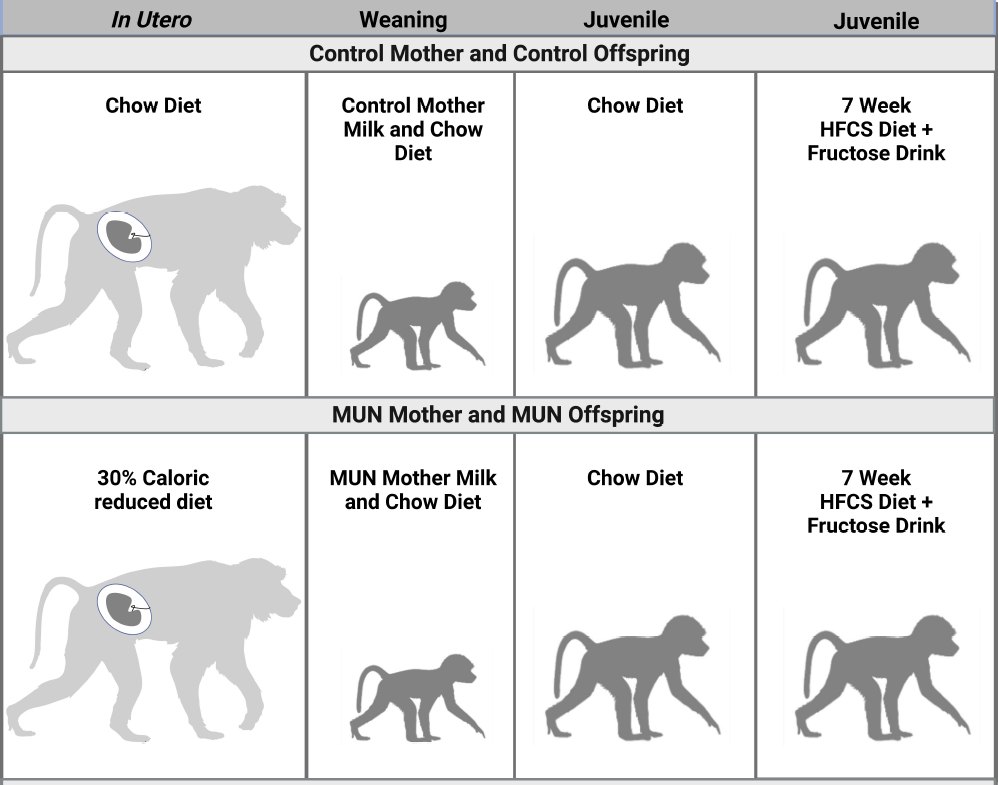
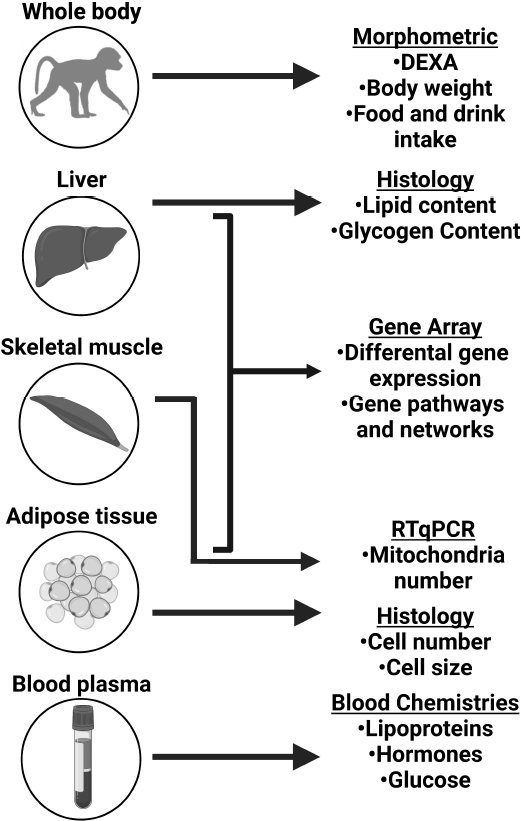
Overview of study design. Offspring investigated in this study were juvenile age from differing maternal *in utero* environments prior to a 7-week high fat, high fructose, high salt diet with high fructose drink (HFCS) (A). Data was collected from whole body, skeletal muscle, liver, adipose, and blood for functional and molecular analysis (B). Created with BioRender.com.

Baboons run once per week into individual feeding cages, passing over an electronic weighing scale, and fed *ad libitum* HFCS diet for 16 hours. High fructose drink and water were provided in Lixit waterers (Lixit, Napa, California) with gauges to measure consumption for each animal, and consumption adjusted to each animals’ body weight.

### Health Assessment and Morphometric Measurements

Animal weights were obtained weekly ^16^. Morphometric measurements collected at baseline and end of challenge, baboons were sedated with 10 mg/kg Ketamine Hydrochloride (Ketaset, Iowa) administered intramuscularly. Hair was removed around the waist and hip circumference lines for accurate measurements. Body length (recumbent length) was measured using an anthropometer (cat. N101, SiberHegner Ltd. Switzerland) from head crown to right tibia. Anterior-posterior abdominal distance was measured from the plane of the back to the anterior point of the abdomen at the level of navel. Waist circumference was measured mid-way between the lowest point of the ribs in the mid-axillary line (costal margin) (10^th^ rib) and iliac crest in the mid-axillary line. Hip circumference measurements at the point of maximum circumference over the buttocks. Body Mass Index (BMI) was calculated by dividing the weight in kilograms by the crown-rump length in cm^2^ (Fig. 1B).

### Blood and Biopsy Collection

Blood and tissue biopsies from liver (right lobe), skeletal muscle (vastus lateralis), and adipose (omental) were collected at baseline and after 7 week challenge. Animals were fasted overnight and immobilized with inhalation anesthesia with isoflurane (1.5%, v/v), and percutaneous venipuncture performed on the femoral vein just caudal to the femoral triangle. A 10 mL vacutainer tube was of serum and 10 mL Na_2_EDTA vacutainer tube was drawn for plasma. Plasma samples were assayed in duplicate. Needle biopsies of 25 mg liver, 5 mg muscle, and 1 gm omental fat were collected from sites aseptically prepared and locally infiltrated with lidocaine. Tissues were placed in tubes, snap frozen in liquid nitrogen, and stored at −80°C.

### Body Composition (DEXA scans)

Body composition was measured by dual-energy X-ray absorptiometry scan (DEXA; Lunar Prodigy; GE Medical Systems, Madison, WI). Animals were placed in supine position on the DEXA bed and extremities positioned within the scanning region. Scans were analyzed using encore2007 software version 11.40.004 (GE Healthcare, Madison, WI). Total body, torso, and each arm and leg, waist, hip region compositions were determined.

### Histological Analysis of Liver and Omental Adipose Biopsies

Frozen fat and liver samples were simultaneously defrosted and fixed in cold formalin, and processed using graded alcohols, xylene, and paraffin embedded. Paraffin blocks were cut in 3 μm sections and processed simultaneously to ensure consistency.

Liver sections were stained with Oil Red O to quantify lipid content (Abcam, Catalog #ab150678) and separate sections with Periodic acid Schiff (PAS) to quantify glycogen content (PAS, Abcam, Catalog #ab150680). Images of sections were captured at 10x magnification using an Olympus BX-41 microscope and QImaging QIcam 12-bit Fast 1394 Camera with Bioquant Osteo 2013 Software. Three slides per animal per diet were analyzed with six pictures (2650 x 1920 pixels) taken from each slide at 2, 4, 6, 8, 10, and 12 o’clock positions and analyzed using NIH Image J software ^18^. Analyses included measuring fraction (area of positive stained x 100%) for Oil Red O, and density (in arbitrary density units) for PAS.

Adipose tissue sections were stained with hematoxylin and eosin, and adipocytes counted using the multipoint tool in Image J. Freehand selection and brush selection tools were used to obtain areas of the field of view without adipocytes. Sums of non-adipocyte areas were subtracted from the total image area to obtain the area of adipocytes. Adipocyte area was divided by total number of adipocytes to obtain average adipocyte size for each field of view. To convert pixel area obtained from ImageJ64 into micrometers, each area was multiplied by pixel size squared (0.769231^2^ = 0.59171633).

### Skeletal Muscle Mitochondrial Number

CDNA was extracted from skeletal muscle biopsies according to manufacturer protocol (Qiagen DNA Purification Kit). Quantitative PCR (qPCR) was performed by SYBR green PCR Kit (Thermo Fisher) with baboon endothelial lipase exon 5 specific primers as the single gene endogenous control (Forward, 5’-TGCACACCCAGGCTTAACTTGT-3’; and Reverse, 5’-CCCAAGACATCGTTGAGTCCAC-3’) and baboon mitochondrial sequence specific primers: (M16SForward, 5’-GCAAACCCTGATGAAGGCTA-3’; and M16SReverse, 5’-GGCCCTGTTCAACTAAGCAC-3’) with 10 ng of DNA per sample run in triplicate, and amplification and quantification by ABI7900 Real Time PCR Instrument. Positive control of a pooled baboon DNA sample and negative no template control were included, each in triplicate.

### Plasma and Serum Measures

Fasting plasma concentrations of glucose (Alfawassermann Cat. # 293598), alanine aminotransferase (ALT; Alfawassermann Cat. # 275454), aspartate aminotransferase (AST; Alfawassermann Cat. # 282242), triglyceride, total cholesterol, LDL, and HDL were determined using an ACE clinical analyzer ^19^. Total hemoglobin (THB) and percent HbA_1c_ were determined from whole blood using an ACE clinical analyzer ^17^. Cortisol was measured by Immulite assay (Siemens/DPC Cat. # 914038), insulin by Immulite assay (Siemens/DPC Cat # 914047), adiponectin by Immulite assay (EMD Millipore HADK1MAG-61K), and leptin by Radioimmunoassay (RIA; EMD Millipore, Cat. # HL-81K).

### Differential Gene Expression

Total RNA was isolated, quality checked, complementary RNA (cRNA) synthesized, and biotinylated as described ^8^ using HumanHT-12 v4 Expression BeadChips (Illumina, Inc.). Gene expression data were extracted and log_2_-transformed using GenomeStudio software (Illumina, Inc.) and analyzed using Partek^®^ Genomics Suite (Partek^®^, St. Louis, MO). Principal Component Analysis (PCA) and hierarchical clustering in Partek^®^ Genomics Suite identified treatment group and diet as the greatest source of variation in each dataset, but no significant contributions by sex for any tissues; therefore, we combined female and male data for analyses. Signal intensities were quality filtered (>0.95), quantile normalized, and differentially expressed genes (DEG) identified by Analysis of Variance (ANOVA; FDR p-value ≤ 0.05, Fold change ≥ 1.2).

### Pathway Enrichment Analysis

Differentially expressed genes (p-value < 0.05) were overlaid onto canonical pathways using Ingenuity Pathway Analysis (IPA; QIAGEN) Knowledge Base. Right-tailed Fisher’s exact test was used to calculate enrichment of DEG in pathways, p < 0.01 ^20^. Pathways with −log p-value ≥ 1.2 and containing DEG FDR p-value ≤ 0.05 were considered significant. Pathways were considered biologically coordinated if gene expression directionality at the end-of-pathway EoP was consistent with overall pathway directionality ^21^.

### Regulatory Network Analysis

Upstream regulatory network analysis was performed with Z-scores predicting regulatory directions and inferring activation/inhibition state of a putative regulator. Detailed statistical models are provided ^22^. Networks were built using the IPA Knowledge Base and required p-values < 0.01, direct connections between molecules based on experimental evidence, network regulator differentially expressed consistent with activation/inhibition status of network, downstream targets differentially expressed, and the network containing at least one FDR DEG (FDR p-value < 0.05) were considered significant.

### Statistical Analyses For Morphometric, Clinical, and Histological Data

As with gene array data, we found no differences between females and males for morphometric, clinical, or histological datasets; therefore, we combined females and males for all analyses. Weight adjusted fructose consumption was calculated by dividing 7-week average fructose consumption by 7-week average body weight. Pairwise comparisons were performed using two tailed t-tests. Data from each diet were analyzed independently using two-way ANOVA. Statistical significance was adjusted for multiple testing with an adjusted p-value < 0.05 considered significant. Regression analysis for fat cell size and adipokine measures were performed using GraphPad Prism (graphpad.com).

## RESULTS

This study included whole animal, hormonal, cellular, and molecular characterization of juvenile MUN offspring of a unique NHP model of developmental programming and matched CON animals (Fig. 1A,1B). Our analysis compared animals before and after a 7-week *ad libitum* HFCS Western style diet supplemented with a high fructose drink (Fig. 1A).

### Comparison of IUGR versus Controls at Baseline Morphometrics

MUN offspring weighed less than CON at birth. At the beginning of the study (baseline), age-matched CON and MUN offspring were the same height; MUN weighed less and had lower BMI (Table 1), suggesting continued impact of MUN during pre-pubertal development. Body composition showed MUN offspring had less fat mass and total mass, and marginally less lean mass than CON (Table 2).

**Table 1:** Weight, height, and BMI measures at baseline and the end of the 7-week challenge

| Group | Weight (Kg) |  |  | Height (cm) |  | BMI (Kg/cm<sup>2</sup>) |  |
| --- | --- | --- | --- | --- | --- | --- | --- |
|  | Newborn | 0wk | 7wk | 0wk | 7wk | 0wk | 7wk |
| CON | 0.919 | 13.91 | 14.77 | 89.61 | 97.13 | 17.34 | 15.64 |
| CON SEM | 0.056 | 0.56 | 0.73 | 1.49 | 2.57 | 0.63 | 0.36 |
| MUN | 0.727 | 11.41 | 12.65 | 88.67 | 93.75 | 14.37 | 14.20 |
| MUN SEM | 0.043 | 1.00 | 1.20 | 2.69 | 2.31 | 0.68 | 0.66 |
| p value 0wk MUN vs CON | **0.022** | **0.024** | 0.166 | 0.734 | 0.370 | **0.001** | 0.085 |
| p value CON 7wk vs 0wk |  |  | **0.040** |  | **0.035** |  | 0.072 |
| p value MUN 7wk vs 0wk |  |  | **0.015** |  | **0.011** |  | 0.344 |

**Table 2:** Body composition from DEXA at baseline and at the end of the 7-week challenge

| Group | bone mass (gm) |  | fat mass (gm) |  | lean mass (Kg) |  | total mass (Kg) |  |
| --- | --- | --- | --- | --- | --- | --- | --- | --- |
|  | 0wk | 7wk | 0wk | 7wk | 0wk | 7wk | 0wk | 7wk |
| CON | 578.3 | 641.4 | 525.4 | 574.7 | 12.29 | 13.13 | 13.40 | 14.35 |
| CON SEM | 29.6 | 38.1 | 22.0 | 26.7 | 0.52 | 0.67 | 0.57 | 0.73 |
| MUN | 468.5 | 550.2 | 406.7 | 479.8 | 9.57 | 11.22 | 10.44 | 12.25 |
| MUN SEM | 56.7 | 69.5 | 40.5 | 52.6 | 0.94 | 1.20 | 1.04 | 1.32 |
| p value 0wk MUN vs CON | 0.101 | 0.256 | **0.021** | 0.120 | **0.023** | 0.175 | **0.025** | 0.175 |
| p value CON 7wk vs 0wk |  | **0.039** |  | **0.024** |  | **0.027** |  | **0.025** |
| p value MUN 7wk vs 0wk |  | **0.012** |  | **0.017** |  | **0.016** |  | **0.0154** |

### Lipoprotein and Hormonal Measures

Plasma measures showed AST, ALT, cortisol, glucose (Table 3), HbA1c, and THB (data not shown) were not different between CON and MUN at baseline. Also, no differences were observed in LDL, HDL, total cholesterol, and triglycerides at baseline (Table 4). Leptin, and adiponectin levels were higher in MUN than CON (Table 5).

**Table 3:**
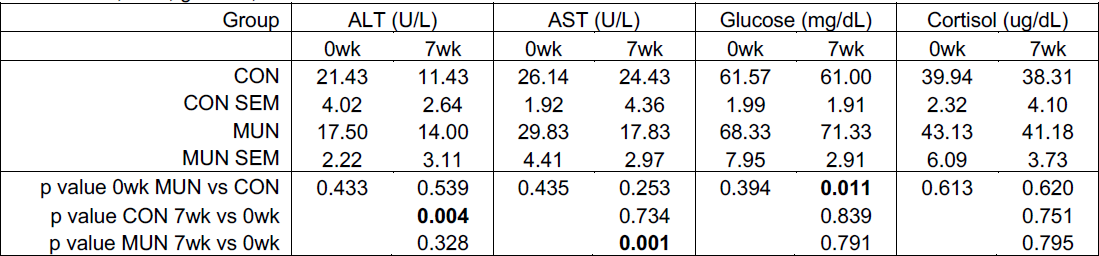
ALT, AST, glucose, and cortisol measures at baseline and at the end of the 7-week challenge (mg/dL)

**Table 4:** Serum lipid measures at baseline and at the end of the 7-week challenge (mg/dL)

| Group | HDL-C (mg/dl) |  | LDL-C (mg/dl) |  | Triglycerides (mg/dl) |  | TSC (mg/dl) |  |
| --- | --- | --- | --- | --- | --- | --- | --- | --- |
|  | 0wk | 7wk | 0wk | 7wk | 0wk | 7wk | 0wk | 7wk |
| CON | 66.21 | 55.21 | 32.29 | 30.29 | 35.86 | 46.29 | 99 | 85.64 |
| CON SEM | 2.55 | 3.37 | 3.31 | 2.78 | 2.82 | 5.19 | 4.12 | 5.16 |
| MUN | 62.92 | 59.92 | 37.17 | 25.5 | 37.67 | 43.83 | 100.33 | 85.58 |
| MUN SEM | 4.33 | 6.77 | 3.77 | 3.53 | 6.97 | 4.85 | 6.36 | 6.89 |
| p value 0wk MUN vs CON | 0.534 | 0.534 | 0.350 | 0.303 | 0.804 | 0.739 | 0.860 | 0.995 |
| p value CON 7wk vs 0wk |  | **0.029** |  | 0.602 |  | **0.039** |  | 0.059 |
| p value MUN 7wk vs 0wk |  | 0.643 |  | **0.015** |  | 0.394 |  | **0.0054** |

**Table 5:** Serum insulin, leptin and adiponectin at baseline and at the end of the 7-week challenge

| Group | Adiponectin (ng/ml) |  | Adipsin (ng/ml) |  | Leptin (ng/ml) |  | Insulin (IU/mL) |  |
| --- | --- | --- | --- | --- | --- | --- | --- | --- |
|  | 0wk | 7wk | 0wk | 7wk | 0wk | 7wk | 0wk | 7wk |
| CON | 53410.9 | 72621.5 | 3658.2 | 5410.0 | 3.12 | 0.96 | 10.33 | 7.98 |
| CON SEM | 5977.1 | 18754.5 | 249.7 | 401.0 | 0.35 | 0.35 | 2.69 | 2.09 |
| MUN | 140429.0 | 85664.6 | 4197.2 | 5860.2 | 5.36 | 2.51 | 31.68 | 14.71 |
| MUN SEM | 36809.1 | 15178.8 | 324.3 | 632.9 | 0.50 | 0.44 | 17.20 | 3.06 |
| p value 0wk MUN vs CON | **0.042** | 0.618 | 0.230 | 0.570 | **0.005** | **0.023** | 0.211 | 0.090 |
| p value CON 7wk vs 0wk |  | 0.280 |  | **0.008** |  | **0.004** |  | 0.522 |
| p value MUN 7wk vs 0wk |  | 0.301 |  | **0.030** |  | **0.020** |  | 0.370 |

### Molecular Analyses of Liver, Skeletal Muscle, and Omental Adipose Tissues

We found the greatest differences between MUN versus CON at baseline in liver with 247 DEG, 194 up-regulated and 53 down-regulated (Table 6, Table S1). Skeletal muscle revealed 84 DEG, 49 up-regulated and 35 down-regulated (Table 6, Table S2). Adipose revealed only one DEG which was up-regulated in MUN compared with CON (Table 6, Table S3).

**Table 6:**
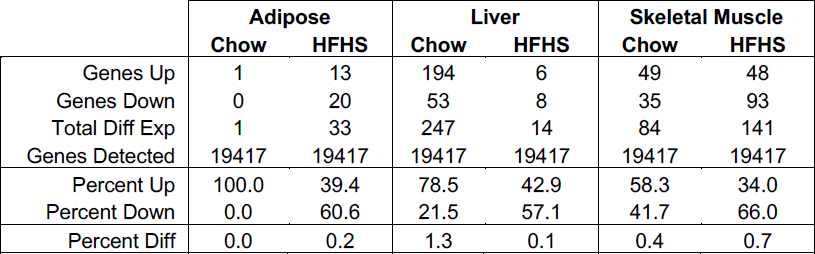
MUN vs CON: Gene Expression Summary FDR corrected p-values, 1.2 FC)

Pathway enrichment analysis reflected DEG numbers with the greatest number of pathways in liver and then skeletal muscle (Table 7). Of the 46 liver pathways, 6 met the EoP criteria; notably, spliceosomal cycle (Fig. S1), unfolded protein response (UPR) (Fig. S2), and BAG2 signaling (chaperone folding activity) (Fig. S3) were up-regulated (Table S4). Skeletal muscle pathways meeting EoP criteria included up-regulation of oxidative phosphorylation (Fig. S4), nucleotide excision repair, and estrogen signaling, with down-regulation of calcium signaling (Fig. S5) and beta adrenergic signaling (Table S5). Significantly different adipose tissue pathways did not include any genes meeting FDR criteria, and of 17 pathways, only ILK signaling met EoP criteria (Table S6).

**Table 7:** MUN vs CON: Pathway Enrichment Summary (includes FDR gene(s) in pathway)

|  | Adipose |  | Liver |  | Skeletal muscle |  |
| --- | --- | --- | --- | --- | --- | --- |
|  | Chow | HFHS | Chow | HFHS | Chow | HFHS |
| Pathways Up | 0 | 1 | 31 | 19 | 3 | 2 |
| Pathways Up EoP | - | 0 | 5 | 0 | 1 | 0 |
| Pathways Down | 0 | 7 | 15 | 1 | 3 | 12 |
| Pathways Down EoP | - | 1 | 1 | 0 | 1 | 0 |
| Total Diff Pathways | 0 | 8 | 46 | 20 | 6 | 14 |
| Percent Up | - | 12.5% | 67.4% | 95.0% | 50.0% | 14.3% |
| Percent Down | - | 87.5% | 32.6% | 5.0% | 50.0% | 85.7% |

Pathway enrichment p-value < 0.01 for DEG p-value < 0.05 and contains FDR gene(s) EoP - pathway met End of Pathway Criteria (see Methods)

Network analysis, similar to pathway analysis, showed the greatest differences in liver with 4 networks (Table 8). One network regulated by KLF6 was inhibited and contained 20 target genes; 75% of the targets’ expression profiles were consistent with network inhibition (Fig. S6, Table S7). Three networks were activated by XBP1 (X-Box Binding Protein 1), a transcription factor that regulates UPR, YAP1 (Yes1 Associated Transcriptional Regulator), a nuclear effector of growth, repair, and homeostasis, and TP63 (Tumor Protein P63), a transcription factor, included overlapping gene targets with a total of 178 gene targets; 25% of the targets’ expression profiles were inconsistent with network activation (Fig. S7, Table S7). Skeletal muscle revealed one network regulated by DDX5 (DEAD-Box Helicase 5), a RNA helicase that coregulates mRNA processing; 10 of the 12 (83%) target gene expression profiles were consistent with network activation (Fig. S8, Table S8) with no networks meeting filtering criteria in adipose (Table S9).

**Table 8:** MUN vs CON: Regulatory Network Summary

|  | Adipose |  | Liver |  | Skeletal muscle |  |
| --- | --- | --- | --- | --- | --- | --- |
|  | Chow | HFHS | Chow | HFHS | Chow | HFHS |
| Networks Up | 0 | 1 | 3 | 3 | 1 | 0 |
| Percent Up-Regulated Genes | - | 53 | 25 | 85 | 83 | - |
| Networks Down | 0 | 2 | 1 | 0 | 0 | 0 |
| Percent Down-Regulated Genes | - | 65 | 25 | - | - | - |
| Total Diff Networks | - | 3 | 4 | 3 | 1 | 0 |
| Percent Networks Up | - | 33.3% | 75.0% | 100.0% | 100.0% | - |
| Percent Networks Down | - | 66.7% | 25.0% | 0.0% | 0.0% | - |

Network p-value < 0.01 for direct connection genes < 0.05 and Network Regulator differentially expressed consistent with network activation/inhibition status

### Cellular Analyses of Liver, Skeletal Muscle, and Omental Adipose Tissues

Due to limited amounts of biopsy material available for cellular analyses of tissues, we focused on phenotypes most relevant to each tissue. Histological analysis of liver showed less glycogen in MUN versus CON (Fig. 2); however, no differences in lipid content were observed (not shown). Histological analysis of adipose tissue showed fat cell size did not differ (Fig. 3), and qPCR of skeletal muscle mitochondria showed no differences between MUN and CON (not shown).

**Fig. 2:**
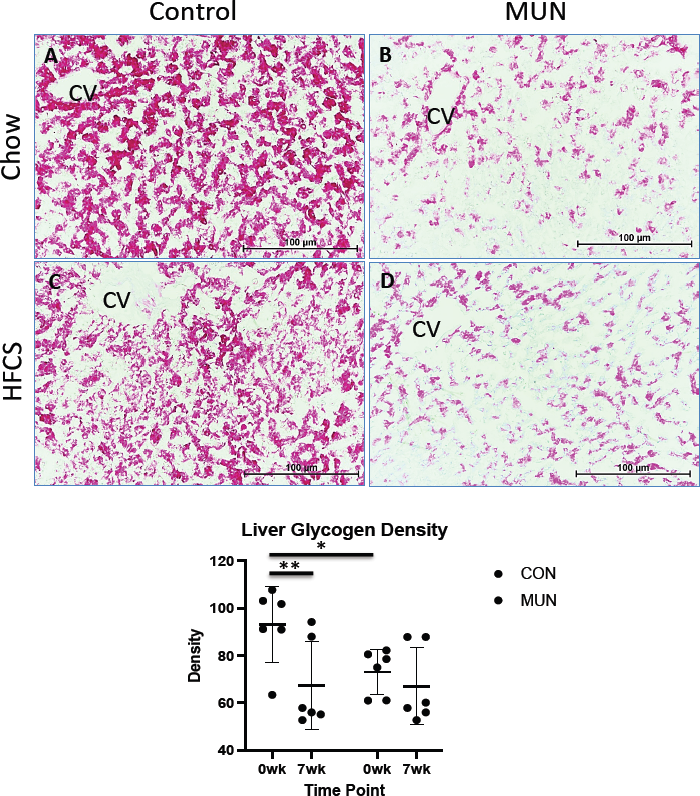
Liver glycogen density from CON (A) and MUN (B) at baseline, CON (C) and MUN (D) after a 7-week high fat, high fructose, high salt diet with high fructose drink. E. shows results for CON (n=7) and MUN (n=6) at both time points. Circles indicate CON and boxes indicate IUGR. * denotes adjusted p < 0.05.

**Fig. 3:**
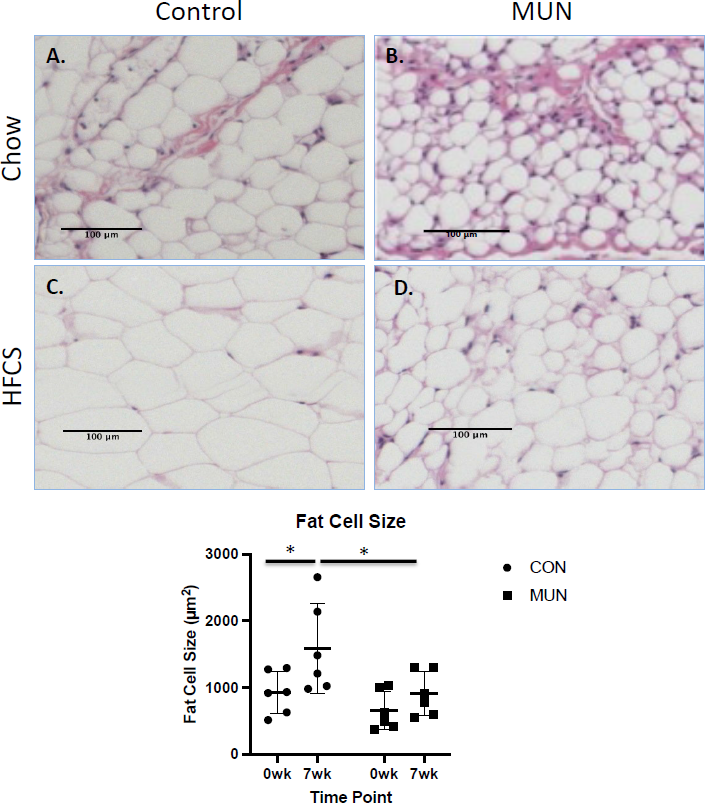
Adipocyte size in representative omental fat sections from CON (A) and MUN (B) at baseline, CON (C) and MUN (D) after a 7-week high fat, high fructose, high salt diet with high fructose drink. E. shows results for CON (n=7) and MUN (n=6) at both time points. Circles indicate CON and boxes indicate MUN. * denotes adjusted p < 0.05.

### Response to HFCS Diet Challenge in IUGR and Controls

To assess whether MUN programs dyslipidemia, diabetes, and appetite in primates, we studied response to a HFCS diet supplemented with a high fructose drink. We monitored drink consumption as an indication of appetitive drive, and quantified changes in body composition and morphometrics, metabolism, liver function, molecular effects, and cellular effects of the challenge.

### Morphometrics and Fructose Drink Consumption

Both groups increased in height from baseline to the end of the 7-week challenge. Although weight and BMI were lower in MUN than CON at baseline, they did not differ at the end of challenge (p ≤ 0.05) (Table 1). In addition, body composition measures of fat mass, lean mass, and total mass did not differ between MUN and CON at the end of the challenge (Table 2), indicating more rapid growth in MUN than CON animals during the challenge. Average sugar drink consumption over 7 weeks, adjusted for each animal’s weight, was approximately 70% greater in MUN versus CON offspring (p < 0.01) (Table 9).

**Table 9:**
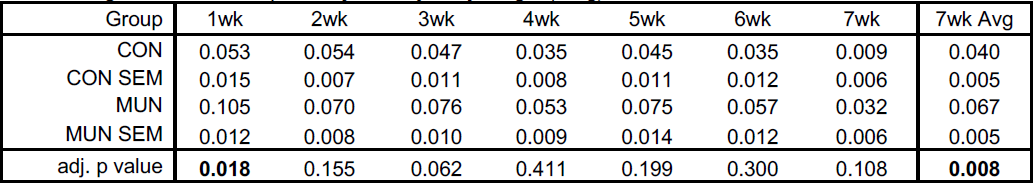
Sugar Drink Consumption adjusted by body weight (L/Kg)

| Group | 1wk | 2wk | 3wk | 4wk | 5wk | 6wk | 7wk | 7wk Avg |
| --- | --- | --- | --- | --- | --- | --- | --- | --- |
| CON | 0.053 | 0.054 | 0.047 | 0.035 | 0.045 | 0.035 | 0.009 | 0.040 |
| CON SEM | 0.015 | 0.007 | 0.011 | 0.008 | 0.011 | 0.012 | 0.006 | 0.005 |
| MUN | 0.105 | 0.070 | 0.076 | 0.053 | 0.075 | 0.057 | 0.032 | 0.067 |
| MUN SEM | 0.012 | 0.008 | 0.010 | 0.009 | 0.014 | 0.012 | 0.006 | 0.005 |
| adj. p value | **0.018** | 0.155 | 0.062 | 0.411 | 0.199 | 0.300 | 0.108 | **0.008** |

### Lipoprotein and Hormonal Measures

Plasma adipsin increased and leptin decreased in both CON and MUN animals in response to HFCS challenge. Cortisol (Table 3), adiponectin, glucose, and insulin did not change (Table 5). Plasma ALT (Table 3) and HDL (Table 4) decreased, and triglycerides (Table 4) increased in CON but not MUN; whereas AST (Table 3), LDL and TSC (Table 4) decreased only in MUN in response to the challenge.

### Liver, Skeletal Muscle, and Adipose Transcriptomes

Transcriptome response to the challenge differed markedly by tissue in MUN versus CON. For CON animals, the greatest response was in skeletal muscle with 332 DEG, 195 up-regulated and 137 down-regulated. In contrast, MUN animals’ greatest response was in liver with 909 DEG, 295 up-regulated and 614 down-regulated. Far fewer DEG were identified in adipose, but it is worth noting that the number of DEG (FDR ≤ 0.05) in CON adipose in response to the HFCS diet was greater than in MUN (CON = 36, IUGR = 8) (Table 10, Supplemental Tables 1 - 3).

**Table 10:** HFCS vs Chow: Gene Expression Summary (FDR corrected p-values, 1.2 FC)

|  | Adipose |  | Liver |  | Skeletal Muscle |  |
| --- | --- | --- | --- | --- | --- | --- |
|  | CON | MUN | CON | MUN | CON | MUN |
| Genes Up | 8 | 1 | 1 | 295 | 195 | 7 |
| Genes Down | 0 | 0 | 8 | 614 | 137 | 27 |
| Total Diff Exp | 8 | 1 | 9 | 909 | 332 | 34 |
| Genes Detected | 19417 | 19417 | 19417 | 19417 | 19417 | 19417 |
| Percent Up | 100.0 | 100.0 | 11.1 | 32.5 | 58.7 | 20.6 |
| Percent Down | 0.0 | 0.0 | 88.9 | 67.5 | 41.3 | 79.4 |
| Percent Diff | 0.0 | 0.0 | 0.1 | 4.7 | 1.7 | 0.2 |

Pathway analysis for CON liver response to HFCS diet showed 8 pathways with 4 meeting EoP criteria, all related to cholesterol biosynthesis and down-regulated (Fig. S9, Table S4), indicating a coordinated response to the HFCS diet in CON animals. In MUN livers, HFCS diet response identified 81 pathways that met FDR DEG requirements, but of these, only one met EoP criteria - Dolichyl-diphosphooligosaccharide biosynthesis - indicating lack of coordinated response to the challenge in livers of MUN animals (Table 11, Table S4). For CON skeletal muscle response, 2 of 24 significant pathways met EoP criteria, oxidative phosphorylation (Fig. S10) and Glycolysis I signaling, both down-regulated. In MUN skeletal muscle HFCS response, 3 of 6 pathways met EoP criteria – oxidative phosphorylation (Fig. S11), calcium signaling, and TCA cycle signaling, all down-regulated. Oxidative phosphorylation down regulation in skeletal muscle was the only pathway common to CON and MUN HFCS response (Table 11, Table S5). In adipose tissue, no pathways met filtering criteria for either CON or MUN (Table 11, Table S6).

**Table 11:**
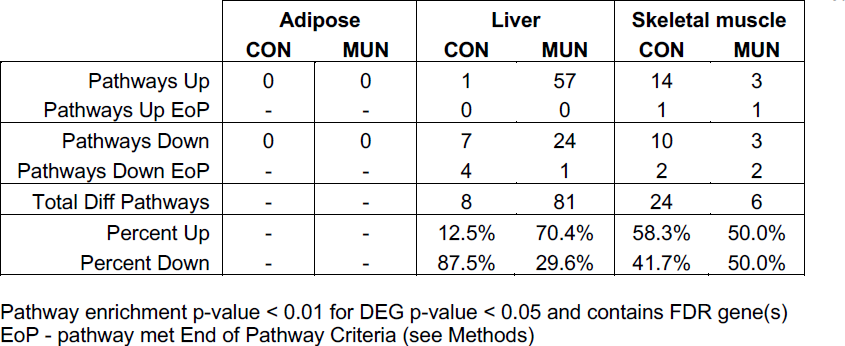
HFCS vs Chow: Pathway Enrichment Summary (includes FDR gene(s) in pathway)

|  | Adipose |  | Liver |  | Skeletal muscle |  |
| --- | --- | --- | --- | --- | --- | --- |
|  | CON | MUN | CON | MUN | CON | MUN |
| Pathways Up | 0 | 0 | 1 | 57 | 14 | 3 |
| Pathways Up EoP | - | - | 0 | 0 | 1 | 1 |
| Pathways Down | 0 | 0 | 7 | 24 | 10 | 3 |
| Pathways Down EoP | - | - | 4 | 1 | 2 | 2 |
| Total Diff Pathways | - | - | 8 | 81 | 24 | 6 |
| Percent Up | - | - | 12.5% | 70.4% | 58.3% | 50.0% |
| Percent Down | - | - | 87.5% | 29.6% | 41.7% | 50.0% |

Pathway enrichment p-value < 0.01 for DEG p-value < 0.05 and contains FDR gene(s) EoP - pathway met End of Pathway Criteria (see Methods)

Network analysis in liver of CON animals’ response to HFCS revealed 7 networks inhibited by NFKBIA (NFKB Inhibitor Alpha), an NFKB inhibitor, FOXO1 (Forkhead Box O1), a transcription factor, STAT3 (Signal Transducer And Activator of Transcription 3), a mediator of cellular responses to growth factors, CEBPB (CCAAT Enhancer Binding Protein Beta), a basic leucine zipper domain containing transcription factor, SREBF1 (Sterol Regulatory Element Binding Transcription Factor 1), a basic helix-loop-helix-leucine zipper transcription factor, NUPR1 (Nuclear Protein 1, Transcriptional Regulator), a transcriptional coactivator, and NR5A2 (Nuclear Receptor Subfamily 5 Group A Member 2), a DNA binding zinc finger transcription factor. These networks included a total of 68 targets with 82% showing expression profiles consistent with inhibitory networks (Fig. S12, Table 12, Table S7). MUN livers showed 4 networks inhibited by XBP1 (X-Box Binding Protein 1), an X-box binding transcription factor, ATF4 (Activating Transcription Factor 4), a transcription factor that binds the cAMP response element, NUPR1 (Nuclear Protein 1, Transcriptional Regulator), a transcriptional coactivator, and MLXIPL (MLX Interacting Protein Like), a basic helix-loop-helix leucine zipper transcription factor that activates carbohydrate response elements in a glucose-dependent manner. The networks for these regulators include a total of 308 targets, with 55% down-regulated (Fig. S13, Table 12, Table S7).

**Table 12:** HFCS vs Chow: Regulatory Network Summary

|  | Adipose |  | Liver |  | Skeletal muscle |  |
| --- | --- | --- | --- | --- | --- | --- |
|  | CON | MUN | CON | MUN | CON | MUN |
| Networks Up | 1 | 1 | 0 | 0 | 2 | 0 |
| Percent Up-Regulated Genes | 61 | 50 | - | - | 60 | - |
| Networks Down | 0 | 1 | 7 | 4 | 2 | 1 |
| Percent Down-Regulated Genes | - | 47 | 82 | 55 | 46 | 98 |
| Total Diff Networks | 1 | 2 | 7 | 4 | 4 | 1 |
| Percent Networks Up | 100.0% | 50.0% | 0.0% | 0.0% | 50.0% | 0.0% |
| Percent Networks Down | 0.0% | 50.0% | 100.0% | 100.0% | 50.0% | 100.0% |

Network p-value < 0.01 for direct connection genes < 0.05 and Network Regulator differentially expressed consistent with network activation/inhibition status

In skeletal muscle, 4 regulatory networks met criteria for CON response to HFCS. Two networks were inhibited: HNF4A (Hepatocyte Nuclear Factor 4 Alpha), a transcription factor, and SMAD7 (SMAD Family Member 7), a signaling antagonist, were activated with 60% of the 221 targets were up-regulated (Fig. S14, Table 12, Table S8); and 2 networks were activated by ESR1 (Estrogen Receptor 1), a ligand activated transcription factor, and IGF2BP1 (Insulin Like Growth Factor 2 mRNA Binding Protein 1), an RNA binding factor that facilitates mRNA transport, but only 46% of the targets were down-regulated (Fig. S15, Table 12, Table S8). For MUN skeletal muscle network response to HFCS, one network regulated by PPARGC1A (PPARG Coactivator 1 Alpha), a transcriptional coactivator, was inhibited with 45 targets and 44 (98%) were down-regulated (Fig. S16, Table 12, Table S8).

In adipose tissue, CON HFCS response revealed one network activated by ESR1 and contained 76 targets of which 61% were up-regulated. This network is of interest in that it contains genes related to wound healing, phagosome formation, glucocorticoid signaling, and G-protein coupled receptor signaling. Also of interest are inclusion of ACACA (Acetyl-CoA Carboxylase Alpha), the rate limiting enzyme in fatty acid synthesis, which has not previously been linked to adipose tissue metabolic signaling (Fig. S17). In MUN adipose tissue, HFCS response revealed one network inhibited by SREBF2 (Sterol Regulatory Element Binding Transcription Factor 2), a transcription factor that regulates cholesterol homeostasis, and one network activated by EPAS1 (Endothelial PAS Domain Protein 1), a transcription factor induced by oxygen. In the SREBF2 network, only 7 of the 15 (47%) target genes were down-regulated (Fig. S18), and for the EPAS1 network, 50% of the 16 target genes were up-regulated. Interestingly, both networks contained genes related to sirtuin signaling, glucocorticoid signaling, TR/RXR activation, and LXR/RXR activation (Fig. S19, Table 12, Table S9). These findings in MUN adipose were similar to liver pathway enrichment in which only one of the 81 pathways met EoP criteria suggesting lack of coordinated molecular response to HFCS challenge in MUN offspring.

### Cellular Response to the HFCS Challenge: Liver, Skeletal Muscle, and Omental Adipose Tissues

Liver glycogen content decreased from baseline to the end of challenge in CON, but not MUN offspring (Fig. 2). Adipocyte size increased in CON from baseline to the end of challenge and was greater than MUN at the end of challenge; while MUN adipocyte size did not change from baseline to the end of challenge (Fig. 3). Interestingly, adiponectin, leptin, and adipsin measures did not correlate with adipocyte cell size for either group on either diet (not shown). Mitochondrial number in skeletal muscle did not differ between MUN and CON (not shown).

## DISCUSSION

A common scenario in many human populations is exposure to a poor nutritional environment *in utero* followed by a postnatal calorie rich environment. This mismatch in available calories is thought to predispose MUN offspring to early onset metabolic dysregulation leading to obesity ^23^, type 2 diabetes ^24, 25^, and cardiovascular disease ^26, 27^. Previous studies have shown that moderately reduced caloric intake in our baboon model detrimentally impacts overall fetal growth leading to IUGR ^28^ with alterations to the fetal heart ^29, 30^, adipose ^31^, kidney ^21, 32, 33^, liver ^10, 34^, and brain frontal cortex ^35^. In addition, IUGR correlates with changes in abundance of appetitive neuropeptides in the ARH of the near-term fetus, which may alter appetitive drive and contribute to obesity in MUN offspring ^9^.

Two major questions from fetal studies are: 1) Do fetal effects of MUN persist in postnatal life of IUGR juvenile offspring? 2) Does a caloric mismatch between *in utero* and postnatal life result in appetitive and metabolic dysregulation in juvenile primates? To our knowledge this is the first study comparing CON and MUN postnatal juvenile NHP, and the impact of total caloric mismatch between fetal and postnatal life on metabolic functions in CON and MUN NHP juveniles.

To address the first question, we compared morphometric measures, appetite, metabolism, and molecular networks in MUN with CON age-matched offspring prior to a HFCS challenge, i.e., animals were maintained on a low cholesterol, low fat chow diet from weaning until the study began.

At baseline, we found fetal programming persisted with MUN offspring weighing less and having less muscle and fat mass than CON. We observed greater serum concentrations of satiety hormones adiponectin and leptin, and lesser amounts of liver glycogen in MUN offspring compared with CON.

Although serum lipid measures, adipocyte size in omental fat, and mitochondrial number in skeletal muscle did not differ between CON and MUN, transcriptome analysis, using stringent filtering criteria, showed marked differences between MUN and CON offspring liver and skeletal muscle. Pathway analysis in liver revealed that few pathways containing DEG had coordinated activity, suggesting molecular dysregulation in many metabolic pathways such as fatty acid beta oxidation, oxidative phosphorylation, and mTOR signaling. It is interesting to note that coordinated pathways of spliceosomal cycling, UPR and BAG2 signaling, which all play roles in molecular diversity, were upregulated in MUN compared with CON – these findings are similar to a previous study by our group showing increased splicing in livers of MUN fetuses ^12^. The lack of coordinated pathways in MUN livers suggests that the potentially increased molecular diversity is not adaptive, but requires further investigation. In skeletal muscle, we observed downregulation of calcium signaling suggesting decreased muscle function in MUN offspring compared with CON. It is worth noting that in a separate imaging study of these animals as young adults, MUN offspring had altered cardiac structure and reduced cardiac function compared with CON ^28^. Based on our findings and the previous study, we provide strong evidence that fetal programming persists in juvenile MUN offspring with overall growth restriction, altered appetitive drive, and dysregulated molecular networks in metabolic tissues (Fig. 4A).

**Fig. 4:**
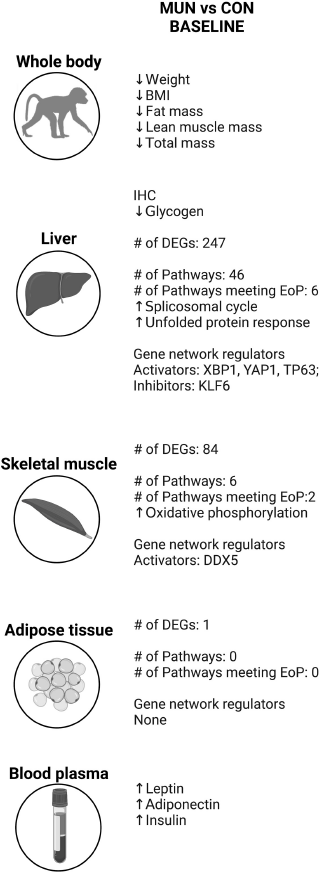
Summary of study findings. Results from comparisons in MUN vs Control animals at baseline (A) are shown with arrows indicating a measure was statistically significantly increased or decreased in MUN animals across whole body, skeletal muscle, liver, adipose, and blood measures. Findings from gene array analysis including gene pathways and networks are also summarized. Results from the 7 week diet challenge compared to the baseline timepoints for CON and IUGR groups respectively follow the same data summary organization as A. Differentially expressed genes (DEGs), end of pathway (EoP). Created with BioRender.com.

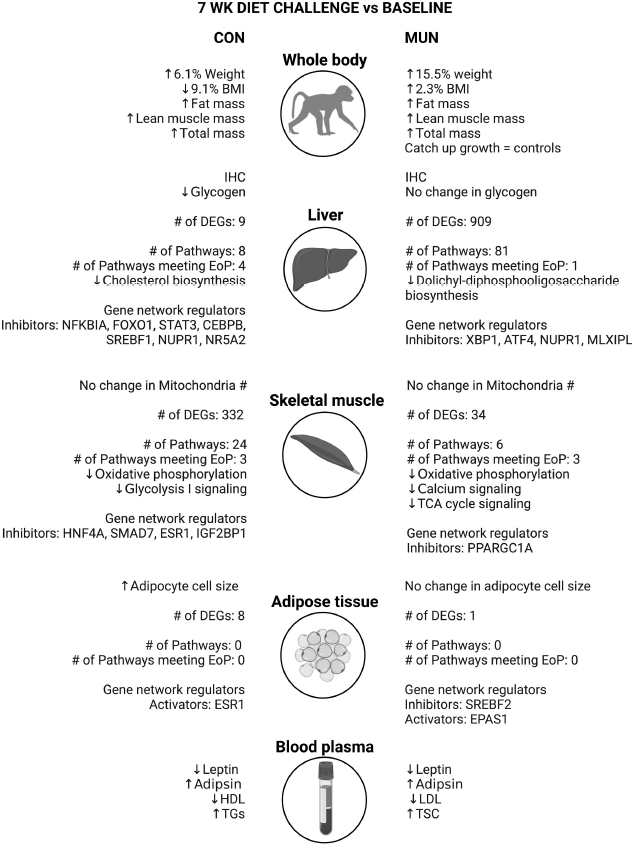

To address the second question of MUN offspring response to caloric mismatch, we compared age-matched CON and MUN juvenile offspring response to a 7-week HFCS diet plus high fructose drink. The challenge aimed to mimic a mismatch of nutrients between fetal and juvenile developmental periods. MUN offspring weighed the same as CON, with no differences in body weight, fat mass, or lean mass at the end of challenge. Few studies have been published evaluating metabolism in mammalian MUN juvenile offspring that demonstrated catch-up growth. One study of IUGR juvenile sheep by Muhle et al. ^36^ found catch up growth in response to an energy dense diet. Studies in humans and animal models have shown that catch-up growth in IUGR offspring is associated with higher abdominal fat mass, blood pressure, total cholesterol, and insulin resistance (e.g., ^37, 38^).

Quantification of satiety hormones showed 2.6 fold greater serum concentrations of leptin in MUN versus CON offspring. These findings are consistent with increased fructose drink consumption in MUN offspring and indicate that satiety signals differ in MUN offspring compared with CON. We did not find differences in measures related to liver function, lipid metabolism, or glucose metabolism. However, we did find differences in liver glycogen content with CON animals showing decreased glycogen content in response to the challenge, with no change in MUN. Our findings are similar to Stanhope and Havel who showed association of failed insulin rise with high fructose consumption and detrimental metabolic consequences such as obesity and insulin resistance ^39^.

It is possible that the effects of MUN on clinical measures related to liver function are not manifest during the juvenile developmental period due to the large metabolic demands of rapid growth. This is supported by the dramatic tissue specific differences in HFCS diet response where CON offspring greatest response is in skeletal muscle, but MUN offspring greatest response is in liver, which is also supported by the differences in liver glycogen with CON animals but not MUN. In addition, CON offspring pathway and network enrichment analyses indicate coordinated molecular response to HFCS challenge, in contrast to a dysregulated response by MUN offspring.

An interesting observation relevant to metabolism and energy management is increased fat cell size in CON with HFCS diet, but not MUN offspring, providing additional cellular evidence of different energy management by MUN offspring than CON, i.e., tissue specific management of excess energy. Previous studies have reported that large adipocytes increase release of inflammatory cytokines with detrimental health outcomes ^40–42^. Although we see an increase in adipocyte size in CON offspring, cell size is still within Category I ^43^. Thus, CON offspring adipocyte size increase may not be metabolically detrimental, but rather a healthy response to a high energy diet in juveniles, i.e., proper storage of excess lipid. In contrast, the lack of adipocyte size increase and greater proportion of small adipocytes in MUN offspring, which has been associated with ectopic lipid accumulation ^44^, suggest a maladaptive response to the high-energy diet. Our findings of fundamental adipocyte differences are supported by molecular data showing coordinated networks in CON but not MUN offspring. In addition, the ESR1 regulated network in CON adipose, includes ACACA, the rate limiting enzyme in fatty acid synthesis ^45^, a gene not previously associated with adipose energy management. Our unbiased molecular analyses may have revealed a novel factor influencing adipocyte function. Taken together, our study reveals fundamental differences between MUN and CON offspring HFCS response at the hormonal, cellular, and molecular levels (Fig. 4B).

## CONCLUSION

The goal of this study was to determine whether *in utero* effects of MUN persist in postnatal juvenile primates, predisposing animals to early onset dyslipidemia and/or metabolic dysregulation. MUN juveniles challenged with an energy dense diet for 7 weeks, a mismatch from the *in utero* environment, demonstrated catch up growth with age-matched CON, significant metabolic differences in hunger and satiety hormones, as well as a non-response of adipocyte size with a high energy diet. Future studies are required to determine whether these alterations manifest as cardiometabolic disease as these animals age.

## SUPPLEMENTARY MATERIALS

For supplementary material for this article, please visit xxx

## DATA SHARING

The gene expression data of this study are openly available in repository Gene Expression Omnibus (GEO Series accession number GSE235713).

## ACKNOWLEDGEMENTS

We gratefully acknowledge the outstanding care by SNPRC veterinary and animal care staff.

## AUTHOR CONTRIBUTIONS

LAC designed and led the study, and drafted the manuscript; KJL, SB, EJD, MJN, CL collected samples and generated data; SP, JC, and AMR contributed to data analysis; PWN, AGC, and MO contributed to study design and interpretation of results. All authors reviewed and edited the manuscript.

## LIMITATIONS

The study had insufficient sample numbers to analyze females and males separately - while there are likely sex-specific differences, we did not have adequate power for such analyses. Animal housing parameters did not allow quantification of food consumption.

---

**Supplemental Figure 1:**
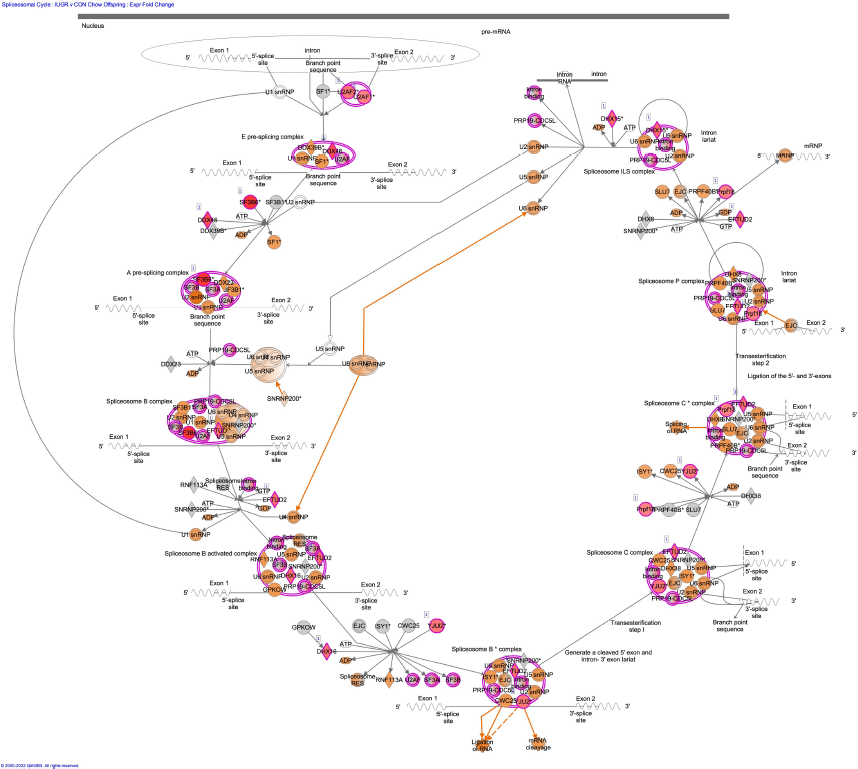
Liver MUN v CON Chow Splicesomal cycle

**Supplemental Figure 2:**
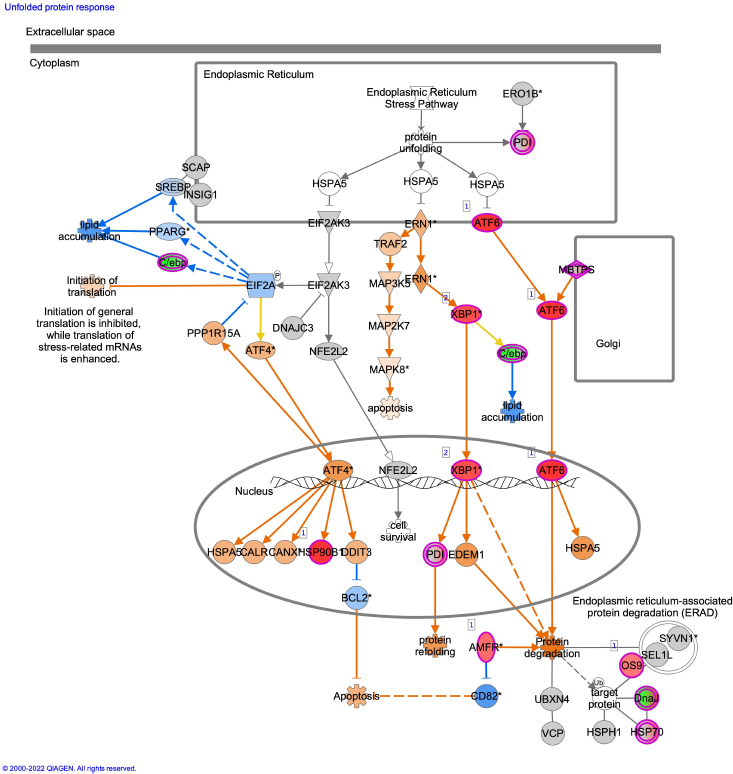
Liver MUN v CON Chow Unfolded Protein Response

**Supplemental Figure 3:**
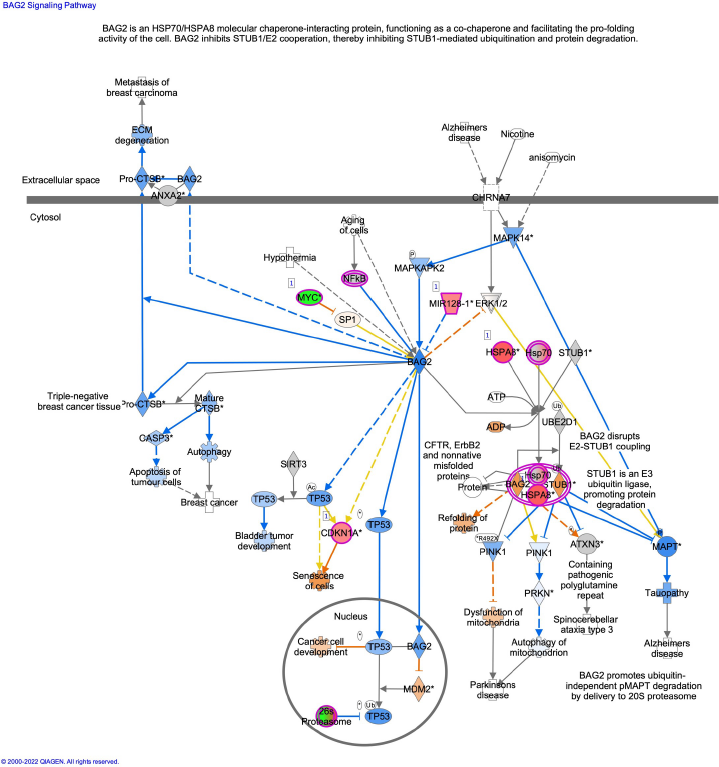
Liver MUN v CON Chow BAG2 Signaling

**Supplemental Figure 4:**
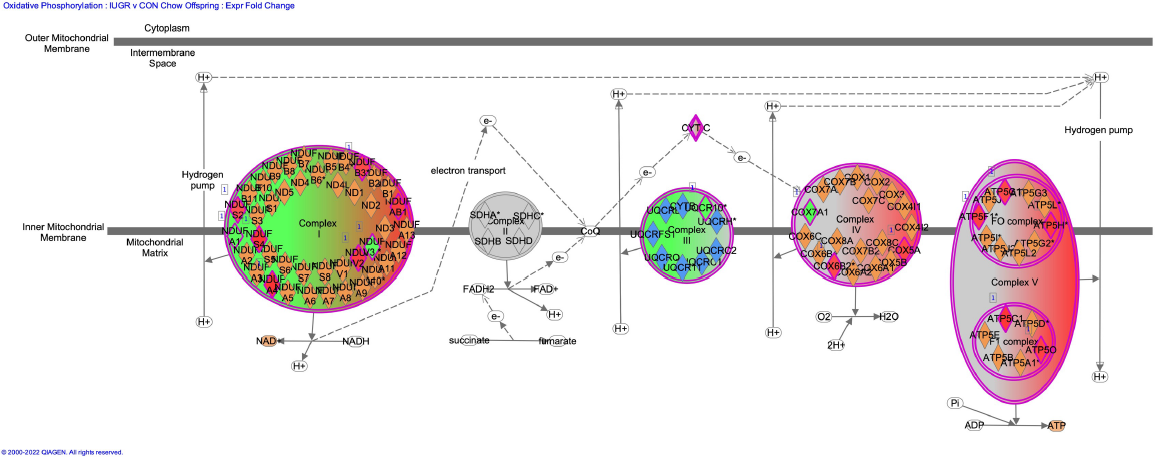
Skeletal Muscle MUN v CON Chow Oxidative Phosphorylation

**Supplemental Figure 5:**
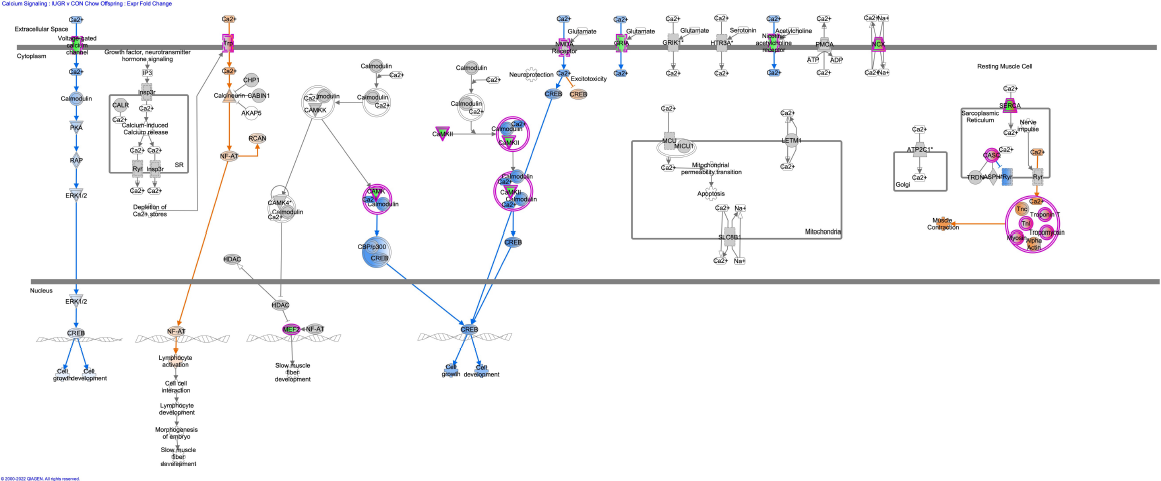
Skeletal Muscle MUN v CON Chow Calcium Signaling

**Supplemental Figure 6:**
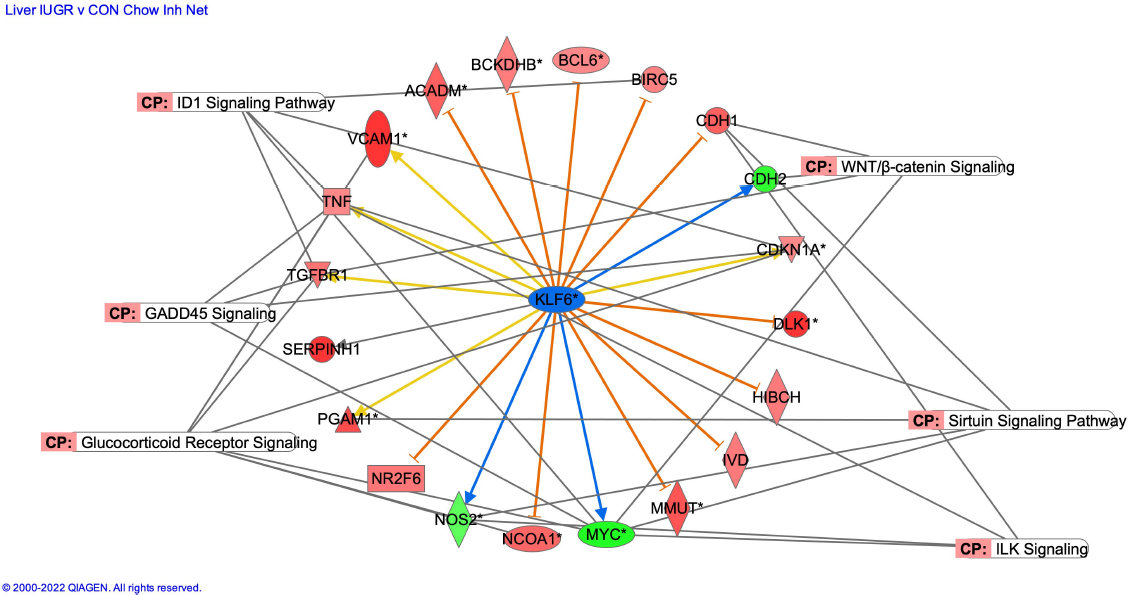
Liver MUN v CON Chow Inhibited Network

**Supplemental Figure 7:**
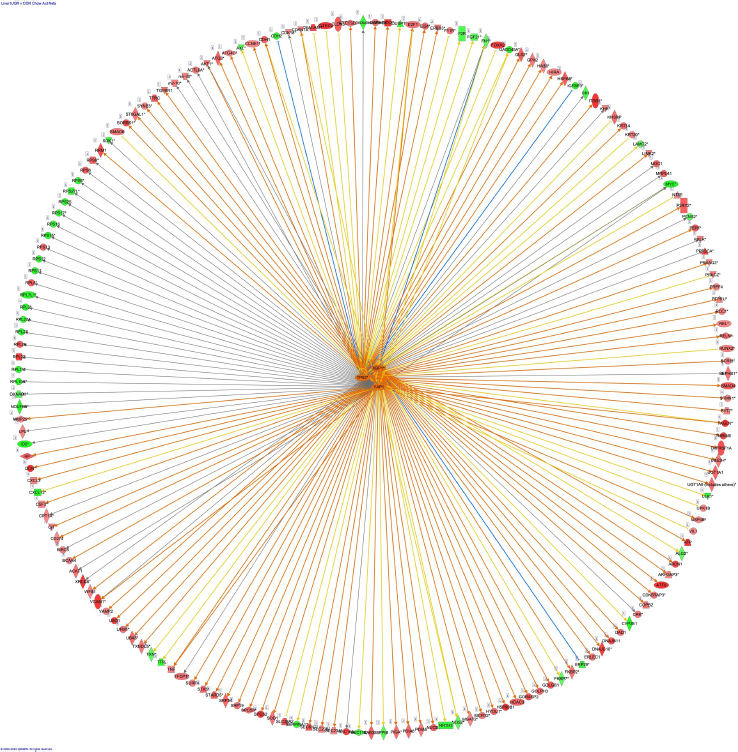
Liver MUN v CON Chow Activated Networks

**Supplemental Figure 8:**
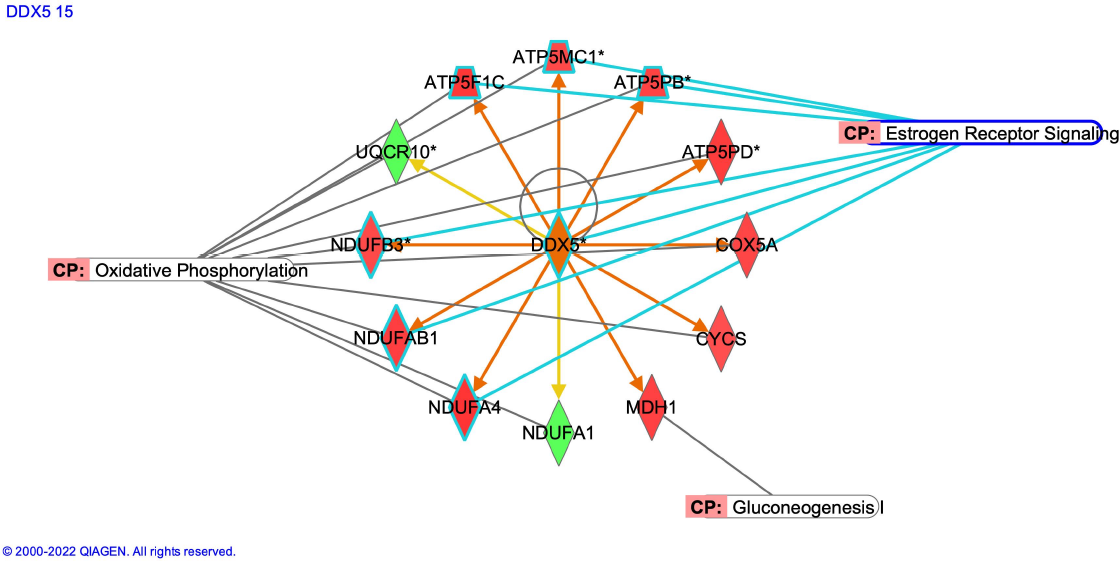
Skeletal Muscle MUN v CON Chow Activated Network

**Supplemental Figure 9:**
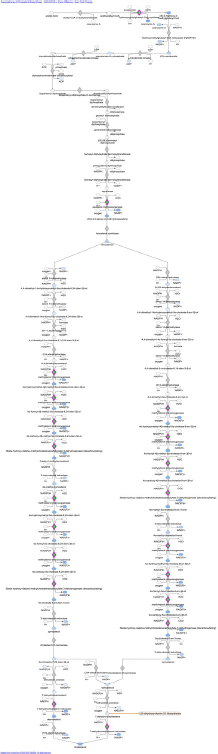
Liver CON HFCS v Chow Cholesterol Biosynthesis

**Supplemental Figure 10:**
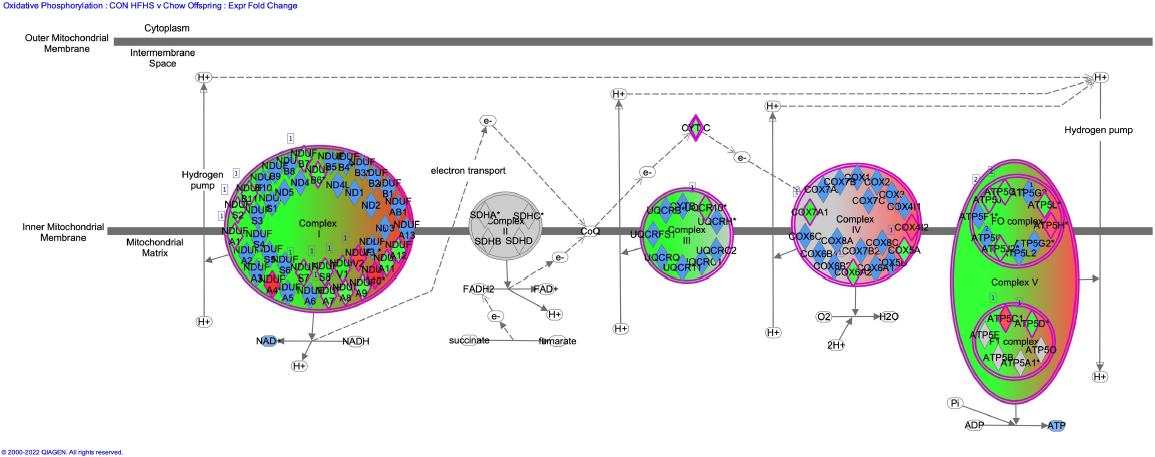
Skeletal Muscle CON HFCS v Chow Oxidative Phosphorylation

**Supplemental Figure 11:**
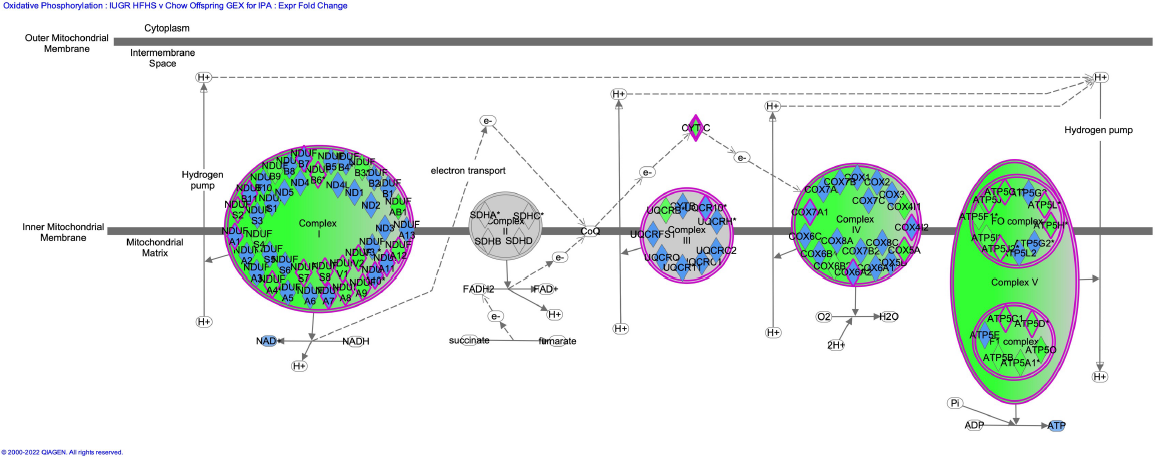
Skeletal Muscle MUN HFCS v Chow Oxidative Phosphorylation

**Supplemental Figure 12:**
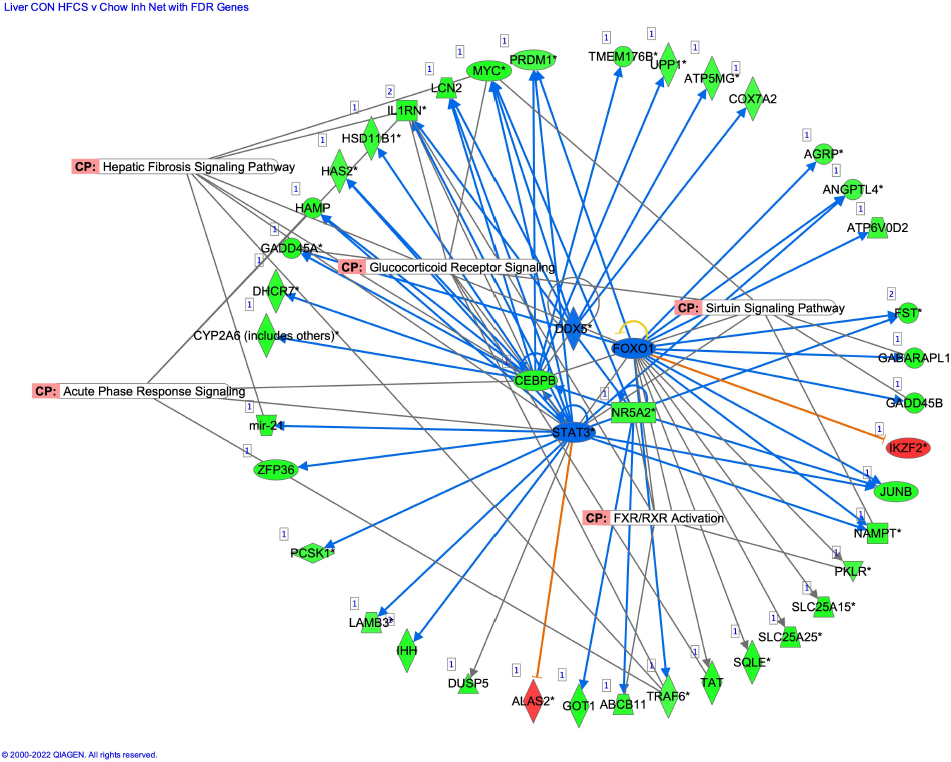
Liver CON HFCS v Chow Inhibited Networks

**Supplemental Figure 13:**
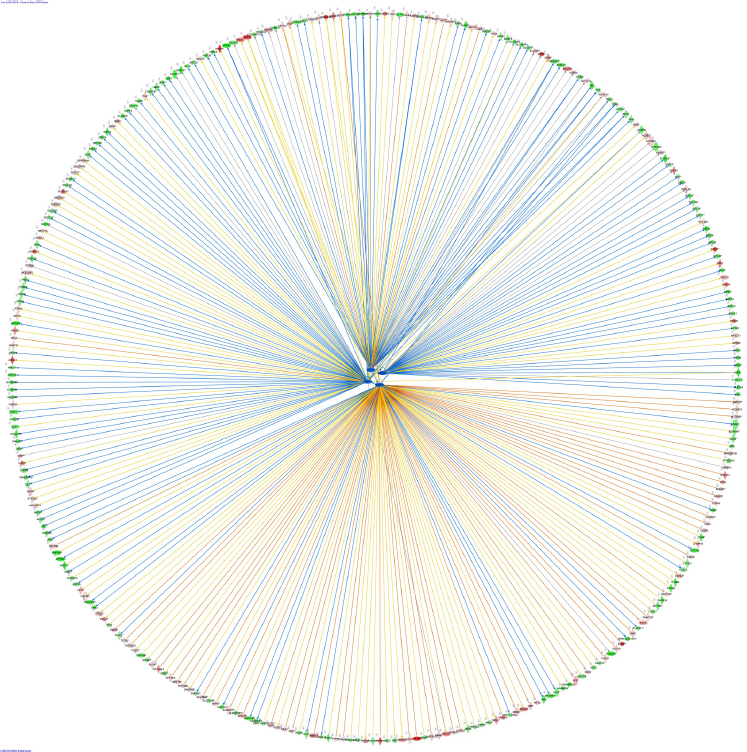
Liver MUN HFCS v Chow Inhibited Networks

**Supplemental Figure 14:**
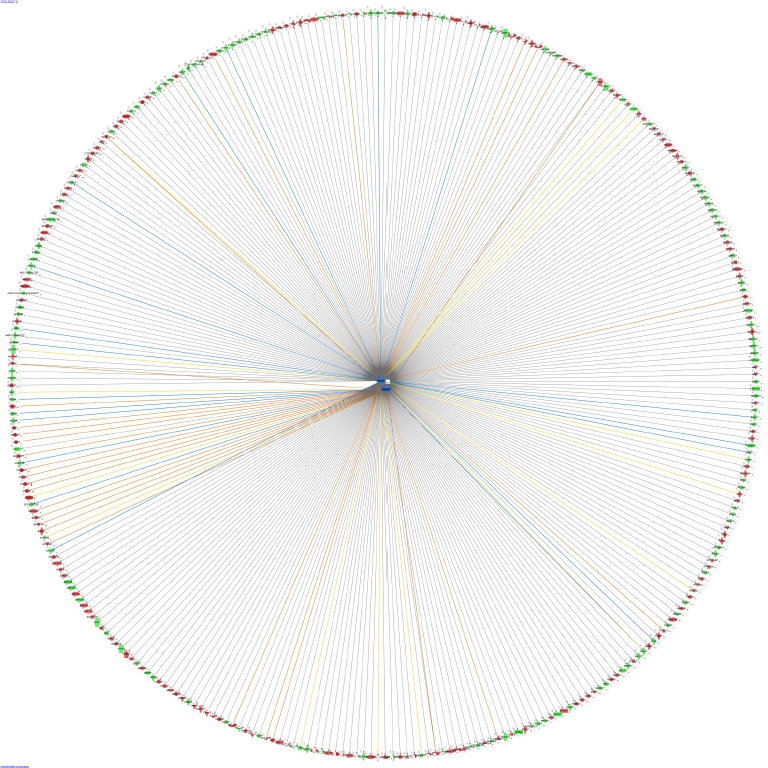
Skeletal Muscle CON HFCS v Chow Inhibited Network

**Supplemental Figure 15:**
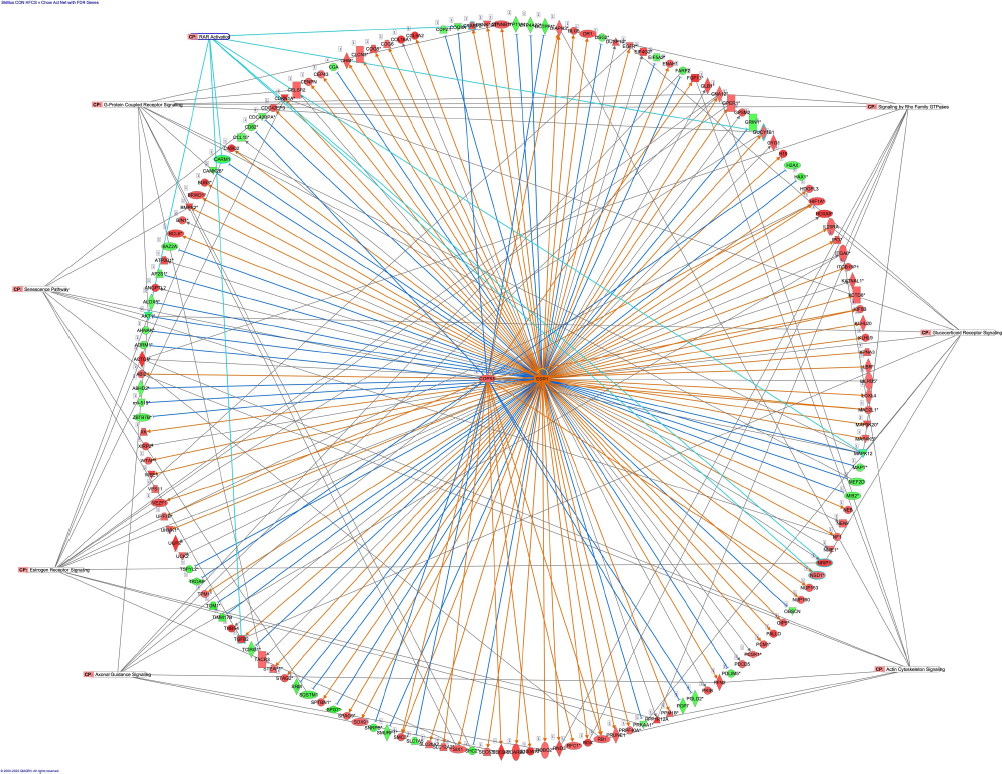
Skeletal Muscle CON HFCS v Chow Activated Networks

**Supplemental Figure 16:**
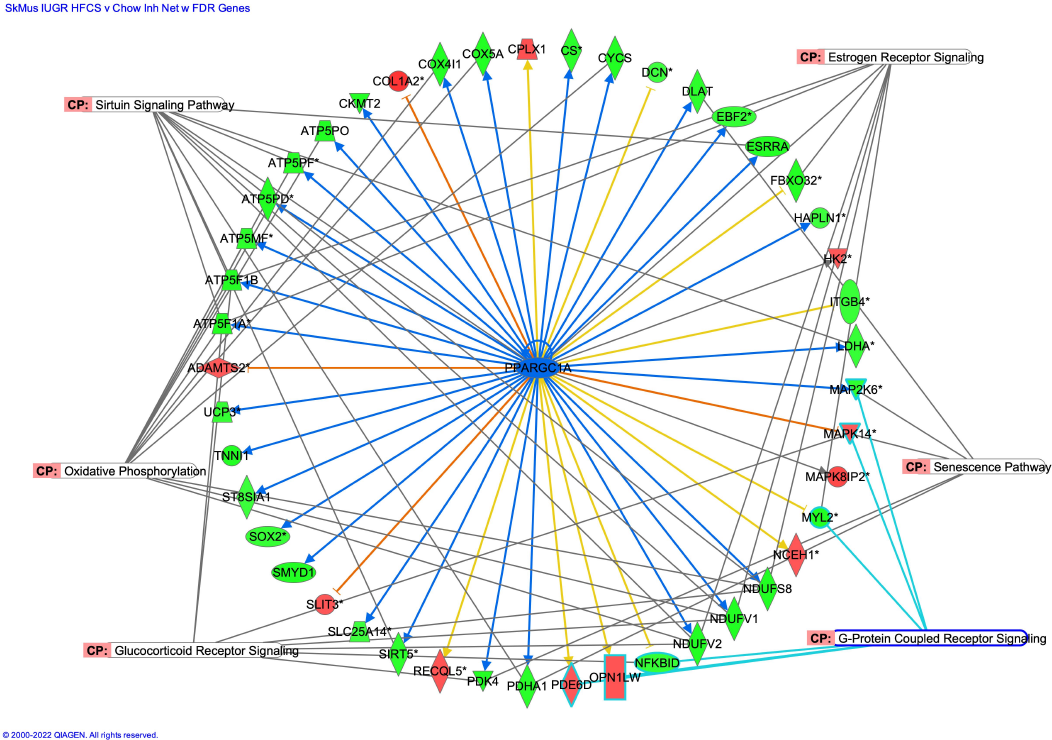
Skeletal Muscle MUN HFCS v Chow Inhibited Network

**Supplemental Figure 17:**
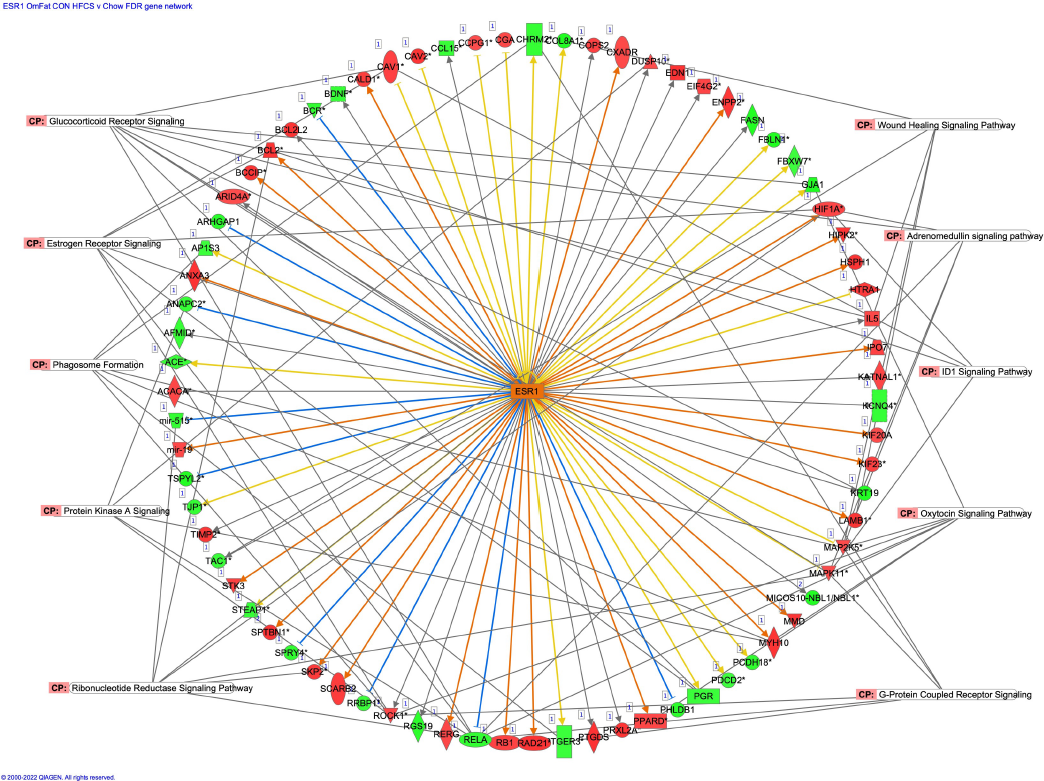
Adipose CON HFCS v Chow Activated Networks

**Supplemental Figure 18:**
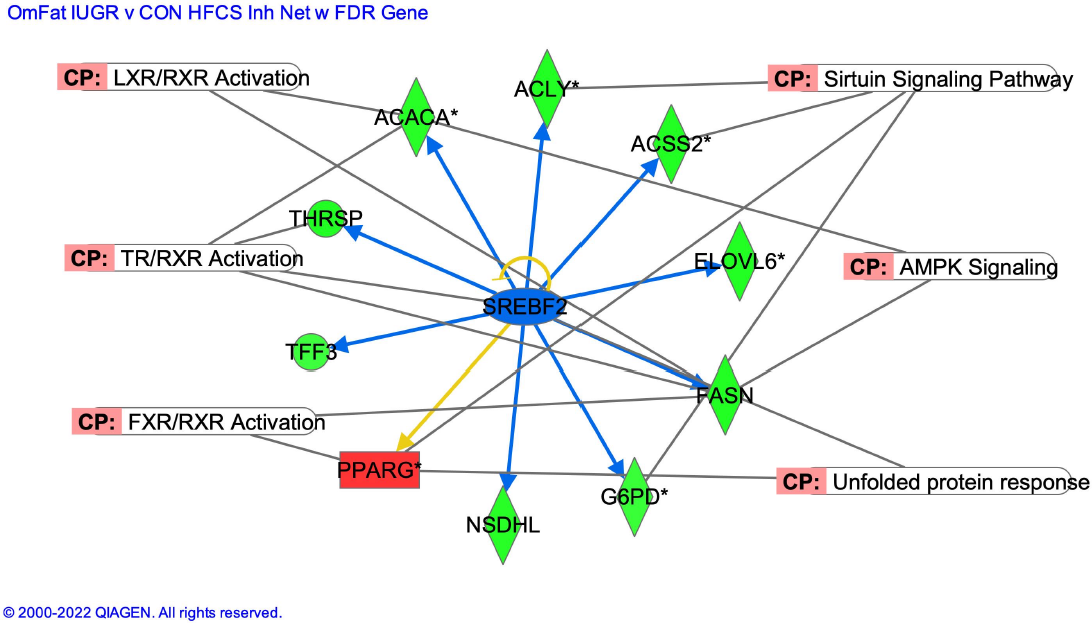
Adipose MUN HFCS v Chow Inhibited Network

**Supplemental Figure 19:**
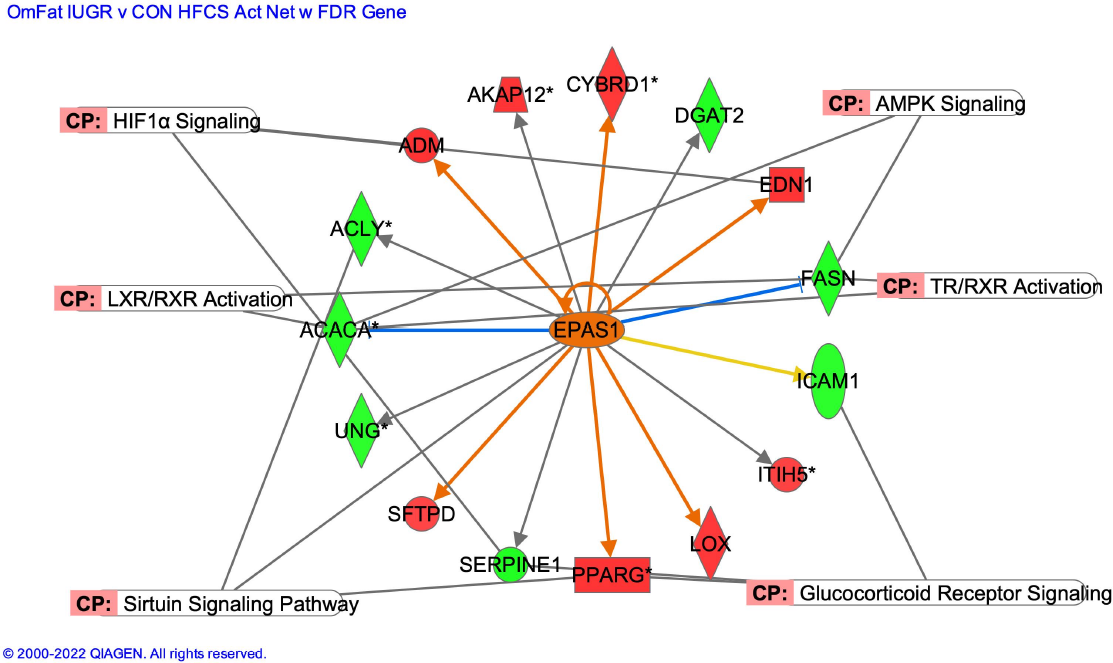
Adipose MUN HFCS v Chow Activated Network

